# Barcoded oligonucleotides ligated on RNA amplified for multiplex and parallel in-situ analyses

**DOI:** 10.1101/281121

**Authors:** Eswar P. R. Iyer, Sukanya Punthambaker, Songlei Liu, Kunal Jindal, Michael Farrell, Jernej Murn, Thomas Ferrante, Stewart Rudnicki, Richie E. Kohman, Asmamaw T. Wassie, Daniel Goodwin, Fei Chen, Shahar Alon, Anubhav Sinha, Denitsa Milanova, Liviu Aron, Conor Camplisson, Alex Skrynnyk, Paul Louis Reginato, Nick Conway, John Aach, Bruce Yankner, Edward S. Boyden, George M. Church

**Affiliations:** Department of Genetics, Harvard Medical School, Boston, MA, USA; Wyss Institute for Biologically Inspired Engineering, Harvard University, Boston, MA, USA; University of California, Riverside, Riverside, CA, USA; MIT Media Lab, Massachusetts Institute of Technology, Cambridge, MA, USA; Department of Biological Engineering, Massachusetts Institute of Technology, Cambridge, MA, USA; McGovern Institute, Massachusetts Institute of Technology, Cambridge, MA, USA; Department of Brain and Cognitive Sciences, Massachusetts Institute of Technology, Cambridge, MA, USA; Division of Health Sciences and Technology, Massachusetts Institute of Technology, Cambridge, MA, USA; Broad Institute, Massachusetts Institute of Technology, Cambridge, MA, USA

**Author notes:** Corresponding author George M. Church.

## Abstract

We present <u>B</u>arcoded <u>O</u>ligonucleotides <u>L</u>igated <u>O</u>n <u>R</u>NA <u>A</u>mplified for <u>M</u>ultiplexed and parallel <u>I</u>n-*S*itu analysis (BOLORAMIS), a reverse-transcription (RT)-free method for spatially-resolved, targeted, in-situ RNA identification of single or multiple targets. For this proof of concept, we have profiled 154 distinct coding and small non-coding transcripts ranging in sizes 18 nucleotides in length and upwards, from over 200, 000 individual human induced pluripotent stem cells (iPSC) and demonstrated compatibility with multiplexed detection, enabled by fluorescent in-situ sequencing. We use BOLORAMIS data to identify differences in spatial localization and cell-to-cell expression heterogeneity. Our results demonstrate BOLORAMIS to be a generalizable toolset for targeted, in-situ detection of coding and small non-coding RNA for single or multiplexed applications.

## Manuscript

Single-cell transcriptomics is an exponentially evolving field, with recent developments in multiplexed in-situ technologies paving the way for spatial imaging of the genome and transcriptome at an unprecedented resolution^1^. We proposed Fluorescent In Situ Sequencing (FISSEQ) in 2003, and in 2014 demonstrated the generation and sequencing of highly multiplexed, spatially resolved in-situ RNA libraries in cells and tissues^2,3^. While FISSEQ’s novelty lay in enabling unbiased discovery, it is primarily limited by poor detection sensitivity, and unsuitability for targeted in-situ transcriptomics (<0.005% compared to single-molecule (sm) FISH)^4,5^. Padlock probes have been demonstrated for in-situ sequencing for multiplexed transcriptomics, with single-base resolution on a small number of transcripts^6^. However in both methods, detection efficiency is a function of RT efficiency, and subject to noise resulting from variable priming efficiency and random priming induced bias^7^. LNA modified primers have been demonstrated to increase RT efficiency with padlock probes (∼30%), but require careful calibration and can be cost-prohibitively expensive for genome-wide applications (∼2 order higher cost than unmodified primers)^5,6,8^.

smFISH based multiplexed transcriptome imaging methods overcome the limitations of RT by directly hybridizing a plurality of oligo-paint like encoded DNA probes directly on target RNA, and subsequently reading out target locations using a two-stage hybridization scheme^9–14^. While smFISH based methods offer the highest in-situ RNA detection efficiency, it is best suited for large transcripts (>1500nt) and can yield lower signal than RCA based methods (∼50-200 vs ∼800-1000 fluorophores/transcript, respectively). As a result, a vast segment of biologically interesting RNA species, (including short-non coding RNA) remain vastly inaccessible to Multiplexed In situ (MIS) methods^1,15^. Consequently, there is a strong demand for robust MIS RNA detection methods that can overcome some of the limitations of each method (sensitivity, cost barrier and transcript-size limitation), while retaining the desirable features of these technologies (scalability, single-nucleotide discrimination, and high detection efficiency).

In order to fill this gap, we developed BOLORAMIS (Barcoded Oligonucleotides Ligated On RNA Amplified for Multiplexed In-Situ), a reverse-transcription free direct RNA detection method for imaging of coding and small-non-coding RNA for multiplexed in-situ applications. BOLORAMIS is based on combinatorial molecular indexing combined with direct RNA dependent ligation and clonal amplification of barcoded padlock probes, and builds on recent improvements in RNA-splinted DNA ligation methods^6,15–23^.

In BOLORAMIS, cells are fixed, permeabilized and incubated with barcoded single-stranded DNA (ssDNA) probes containing two target-RNA complementary termini and a linker segment containing a universal sequencing anchor. The sequencing anchor is flanked by dual 6-base barcodes to allow sufficient barcode-diversity for whole-transcriptome multiplexing (∼10^6^ unique barcodes). The oligonucleotides are directly ligated on target RNA with PBCV-1 DNA ligase to generate circularized barcoded ssDNA probes. Single-molecule signal is then amplified with rolling circle amplification (RCA) with the incorporation of aminoallyl deoxyuridine 5′-triphosphate (dUTP). The amplicons are crosslinked to the cellular protein matrix, using an amine-reactive bifunctional linker to generate spatially structured three-dimensional FISSEQ libraries (Figure 1A)^3,5^.

**Figure 1:**
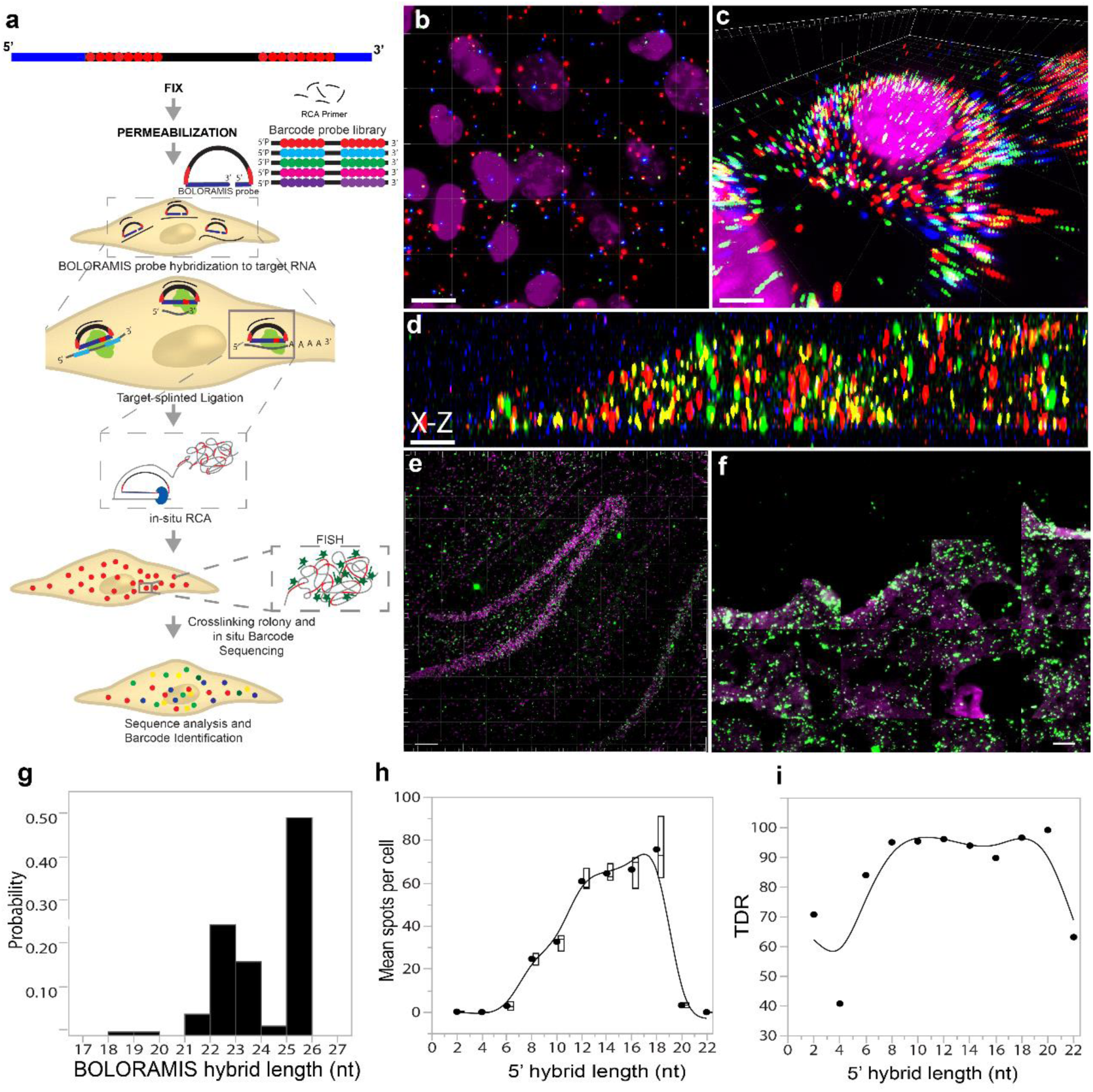
In-situ BOLORAMIS RNA detection on cells and tissues: (a) Outline of BOLORAMIS strategy b) demonstration of multiplexed in-situ miRNA library (77-plex), with a single base sequence in 3-color space (scale bar=5 µm). c) 3-dimensional rendering of a single MEF7 cell showing bright-fluorescent BOLORAMIS spots from multiplexed mRNA targeting. Z-slices are seen as stacked signal in in 3-D rendering (scale bar=7 µm). d) A magnified in-situ sequenced X-Z rendering of a single HeLa cell showing bright, spatially-resolved clonal BOLORAMIS amplicons. Z-axis artifact is seen as elongation of puncta by 3-4-fold diameter (scale bar=5 µm). Each color corresponds to a unique sequence in a barcode space. e) BOLORAMIS capture of single GAPDH mRNA molecules in frozen mouse (e) and human (f) brain sections. BOLORAMIS amplicons were visualized using a FISH probe targeting a universal probe sequence. BOLORAMIS detection of GAPDH in frozen mouse brain section f) zoomed in-view of mouse brain showing single GAPDH RNA locations (green) and nucleus (magenta). g) Distribution of BOLORAMIS RNA: DNA hybrid length for 269 probes. h) BOLORAMIS relative detection sensitivity as a function of 5’ hybridization sequence length. i) BOLORAMIS TDR as a function of 5’ hybridization sequence length. Scale bars: 20, 7, 3, 50 and 50 µm for b, c, d, e and f respectively.

Using BOLORAMIS, we were able to readily generate dense punctate amplicons from targeted RNA detection in a wide range of continuous cell lines, as well as in frozen human and mouse brain sections (Figures 1, S1-S3). Negative control probes (non-targeting), or absence of PBCV 1 DNA ligase resulted in almost no signal as expected (S1-f, g). We further confirmed detection specificity by designing a probe targeting human 3’ untranslated region (3’UTR) of Actin B gene (ACTB) with ∼95% sequence identity to mouse ortholog, and tested detection on both mouse and human cell-lines. BOLORAMIS specifically generated dense amplicons in human HeLa cells, but not in mouse NIH-3T3 fibroblasts (figure S1, supplemental table S9). As control, targeting a perfectly conserved 3’ UTR sequence resulted in comparable number of amplicons in both human and mouse cell lines (figure S2).

Next, we wanted to characterize BOLORAMIS relative detection sensitivity and true discovery rate (TDR) as a function of the length of the 5’ targeting arm length for a probe with a total conserved RNA hybrid sequence length of 25 nt. We systematically tested probes with a range of 5’ targeting hybrid length (from 2 to 24 bases) and quantified the observed mean spots per cell for each condition for *ACTB*. False-discovery rate (FDR) rate was estimated by a mismatch probe containing a single base mismatch at the 3’ terminus for each condition. To minimize bias, the donor/acceptor ligation junction bases were kept constant in all 5’/3’ RNA hybrid length combinations. Overall, we observed an asymmetric increase in relative detection sensitivity with increasing 5’ arm length, peaking at position 18 with a TDR of 96.6%, with ∼29 fold-discrimination between probes with a single base 3’ mismatch (Figure 1-h, i, supplemental figure 1, 3, supplemental table 1). These results indicate that this method is suitable for direct-RNA detection with high specificity, nearing single-base resolution (supplemental S3).

We next evaluated BOLORAMIS performance in detecting a wide range of coding and non-coding RNA targets in a biologically meaningful context. We designed 269 probes targeting 77 miRNA and an equal number of Transcription Factors (TF’s), expressed at varying abundance levels in human iPSC cells (supplemental methods, supplemental table 7)^24^. Each probe was tested individually in-situ in PGP1 human iPSC cells in 384-well plates. Single-cell spot counts were quantified from a total of 217,206 human iPSC cells using automated imaging and image-processing pipeline (Figure 2, supplemental figure S4, Supplemental Pipeline, Supplemental table S9).

**Figure 2:**
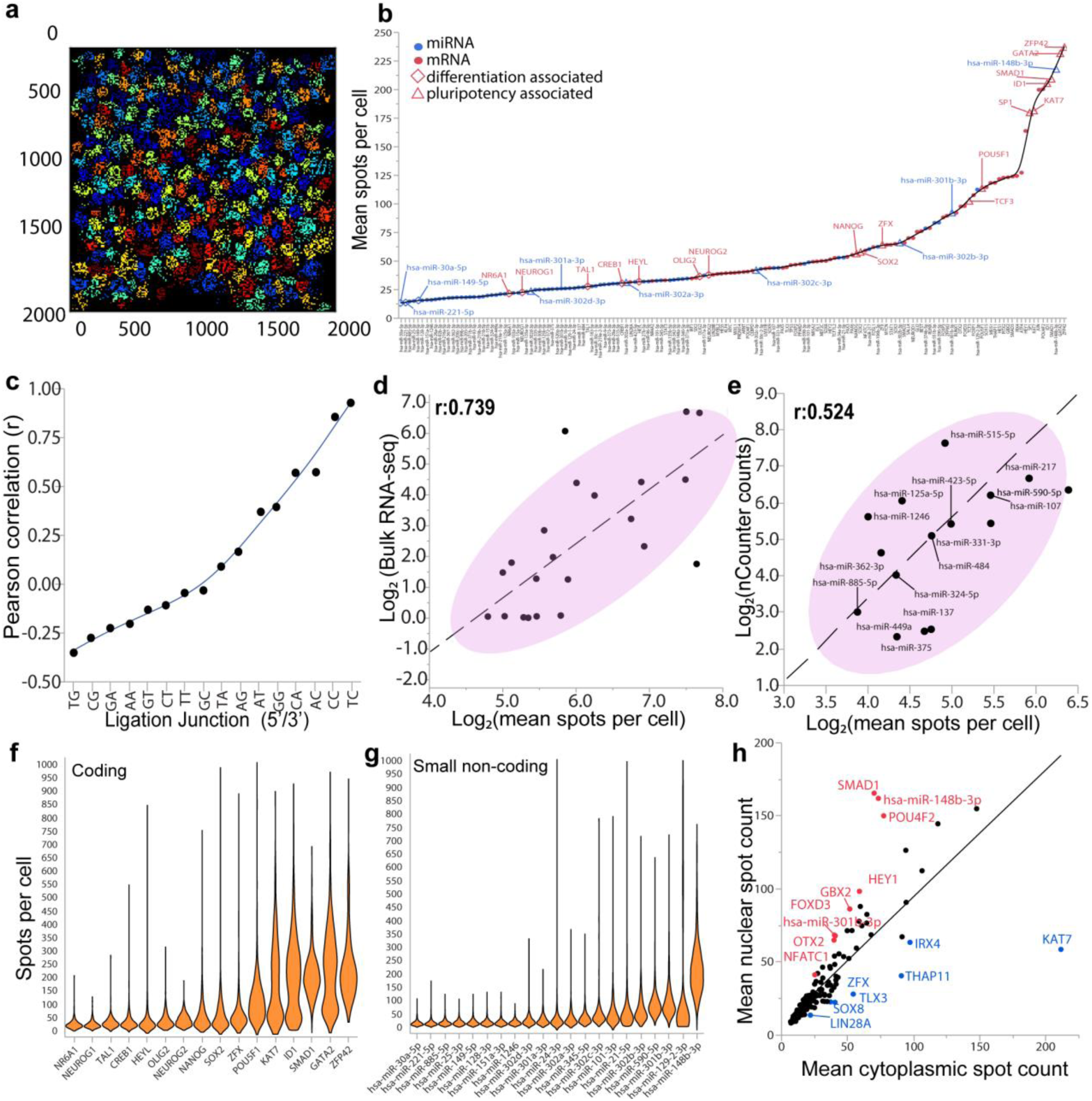
Quantitative BOLORAMIS detection of coding and non-coding RNA: a) BOLORAMIS single-cell expression values are collected from automated segmentation, which involves digitally assigning each spot to a single cell. Spots assigned to different cells are shown in distinct colors. Image of automated segmentation shows BOLORAMIS spots assigned to single cells. b) Mean BOLORAMIS spot/cell measurements for 154 mRNA and miRNA measured from human iPSC cells. c) Effect of ligation junction composition on Pearson’s correlation of mRNA measurements with bulk RNA-seq d) Comparing BOLORAMIS values from probes with top three ligation junctions (TC, CC, AC) with bulk RNA-seq resulted in a Pearson’s correlation coefficient of 0.739. e) Comparison of BOLORAMIS miRNA measurements with previously published bulk nCounter profiling from hiPSC. f, g) Single cell BOLORAMIS expression distribution of key coding and small-noncoding cell-type markers are shown. h) Scatterplot showing nuclear to cytoplasmic RNA distribution. N: C ratio values with Z scores>=/<= 1.64 are highlighted in red and blue colors respectively.

**Figure 3:**
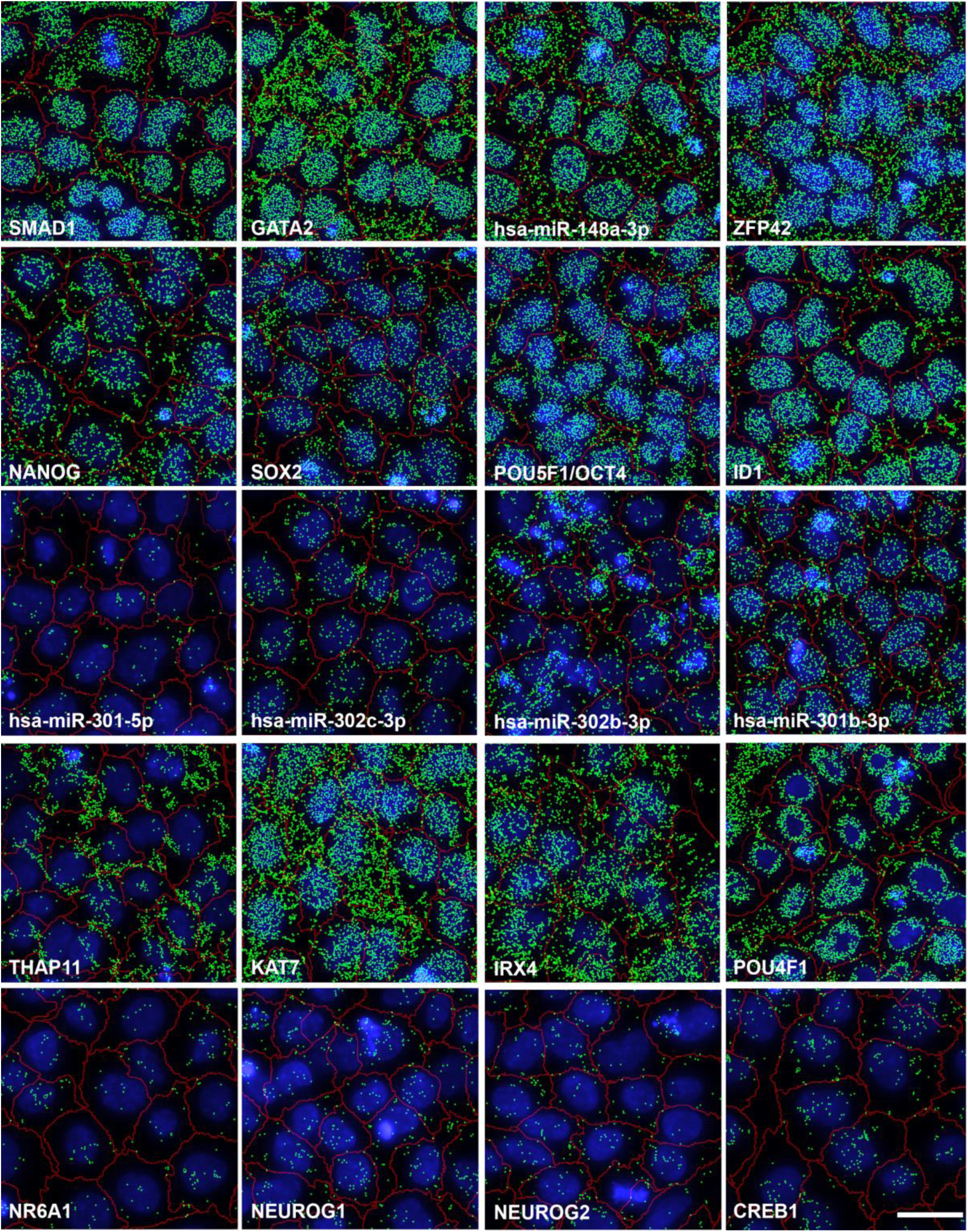
Examples of BOLORAMIS microscopy phenotypes of coding and small-non coding RNA. CellProfiler **s**egmented cell boundaries are indicated as red outlines. Detected spots are indicated by green diamonds for clarity. Scale bar: 20 µm

On a whole, BOLORAMIS expression values were indicative of a stem cell expression signature (Figure 2, b, f, g). For example, some of the highest ranked mRNA corresponded to pluripotency markers (*ZFP42, GATA2, SMAD1, ID1, KAT7, OCT4, SOX2, NANOG* and *ZFX*) (Figure 2, b). Conversely, mRNA with the lowest BOLORAMIS single-cell expression values were associated with promoting cellular differentiation. For example, the lowest ranked TFs were *NR61A, NEUROG1, LIN28B, TAL1, NR4A2, CREB1, OLIG2* and *NEUROG2* (Figure 2, b). Single cell BOLORAMIS expression counts varied between 1-1000 spots/cell with a mean of 53.8 +/-74.9 Stdev. (Supplemental Figure S5). The highest detected mean BOLORAMIS expression value was 236.95 puncta/cell (*ZFP42*, a pluripotency gene), which is comparable with the reported upper-limit of high-throughput bDNA sm-FISH detection at similar magnification (Figure 2, b, e)^12,25^. For a given probe, we observed highly reproducible mean spot counts per cell for independent BOLORAMIS assays (Pearson’s r: 0.859, Supplemental Figure S6, a). We estimate BOLORAMIS to perform better than RNA-seq for transcripts expressed at low abundance (<=5 FPKM). On average, we observed over 71 spots per cell with BOLORAMIS for transcripts with bulk RNAseq FPKM values <=5 (which is roughly estimated to be equivalent to ∼1 transcript per cell)^26^.

We next compared BOLORAMIS mRNA measurements with published RNAseq values. As a population, BOLORAMIS probes exhibited a low-positive correlation with bulk RNAseq values (Pearson’s r: 0.142). We suspected that the low correlation might likely be resulting from poor ligation-efficiencies for certain donor/acceptor pairs^23^. To test this hypothesis, we split our data from 192 probes into 16 donor/acceptor ligation junction categories, and found a close dependency between Pearson correlation and ligation junction composition, which ranged from −0.351 to 0.928 across all 16 categories. Consistent with previous reports, probes with a G in either donor or acceptor positions correlated poorly with bulk RNAseq measurements (Figure 2, c)^23^. Interestingly, C in donor/acceptor positions (dT/C, dC/C, dA/C or dC/A) displayed the highest correlation with RNAseq values (Pearson’s r: 0.928, 0.856, 0.573 and 0.57 respectively) (Figure 2, c). Probes with best ligation junctions (TC, CC and AC) display in a higher Pearson’s correlation with bulk RNA-seq (0.739, Figure 2, d).

We also observed a dependency between probe melting temperatures (T_m_) and Pearson’s correlation with bulk RNAseq values, which explained some of the variance associate with the data. In general, probes with greater Tm’s displayed a higher Pearson’s correlation with bulk RNAseq values (R^2^ = 0.721, supplemental table S11). Due to RNA size constrains, miRNA, targeting probes were designed to avoid G in either donor or acceptor position to improve ligation efficiency, and exhibited a Pearson’s correlation of 0.524 with published bulk miRNA nCounter expression measurements in PGP1 hiPSC (Figure 2, e and supplemental table 3)^24^.

Single-cell expression measurements allow unique insights into population-heterogeneity, which is not possible with bulk methods that average population signal. To assess single-cell expression variability over a population of cells, we calculated the coefficient of variation (COV), defined as the ratio of standard deviation to mean, expressed in percentage for all targets. The mean COV of expression for all targets was 77.3 +/-27% (Stdev) and ranged from 38.7% COV to 217.1% COV (Figure 2-f, g). Independent probes targeting the same mRNA, on average displayed a lower variability than the observed overall single-cell expression heterogeneity. The mean COV between at least two independent probes targeting the same RNA was 52.63 +/-29.46% (Stdev, n=53, Supplemental Figure S4-b).

Single-cell spatial distribution of BOLORAMIS spots seemed non-random, and consistent for a given probe. To quantify differences in spatial localization, we calculated the ratio of mean nuclear to cytoplasmic puncta counts (N: C ratio) and identified several transcripts with signal preferentially localized in the nucleus or cytoplasm in iPSCs (Figure 2, h). For example, we observed a significantly higher nuclear signal from probes targeting SMAD1, hsa-miR-148b-3p, POU4F2, HEY1, GBX2, FOXD3, hsa-miR-301-3p, OTX2 and NFATC1 (Z-score N: C ratio >=1.64) (Figure 2, h). In contrast, probes targeting KAT7, IRX4, THAP11, ZFX, TLX3, SOX8 and LIN28 showed higher cytoplasmic localization (Z-score N: C ratio <=-1.64) (Figure 2,h, supplemental Figure S6).

We tested our ability to generate multiplexed in-situ targeted RNA detection libraries by pooling individual BOLORAMIS probes targeting either miRNA (77 plex), mRNA (77 plex) or splice-junctions (18 plex) in both HeLa and hiPSC cell lines. To verify compatibility with in-situ sequencing chemistries, which depends on clonal, spatially resolved fluorescent amplicons, we quality-checked all our libraries with single-base sequencing, which resulted in bright, punctate spectrally resolved fluorescent spots in the expected color-space (Figure 1, b, c, d, Supplemental Figure S6). BOLORAMIS amplicons displayed high stability, and withstood multiple imaging and sequencing cycles (Supplemental Figure S8).

As a proof of concept for simultaneous multiplexed detection of both coding and small non-coding RNA with FISSEQ, we performed a pooled BOLORAMIS library construction with probes targeting 14 transcripts, consisting of 4 miRNA and 10 mRNA. We sequenced 3 bases of the barcodes using sequence by ligation chemistry and extracted the relative barcode frequencies with 98.26% accuracy (511 correct barcodes out of 520, Figure 4). Barcode enrichment was non-random, and only 14 of 64 possible triplet barcodes were observed in our data, of which 9 were expected within our library. Barcodes with highest frequencies corresponded to iPSC markers ID1, NANOG, hsa-miR-148b-3p, Sox2, GATA and KAT7 (Figure 4, b and Supplemental table 7). Interestingly, we did not detect barcodes corresponding to ZEPF42 and POU5F1 in our data. We believe this is likely a barcode mis-representation artifact resulting from single-base mis-reads during base-calling. Several ongoing developments in our lab, and others should address these in the near future with the use of error-correction barcodes, longer sequencing reads, better sequencing chemistries and improved base-calling algorithms (Supplemental Figure S8)^27,28^.

**Figure 4:**
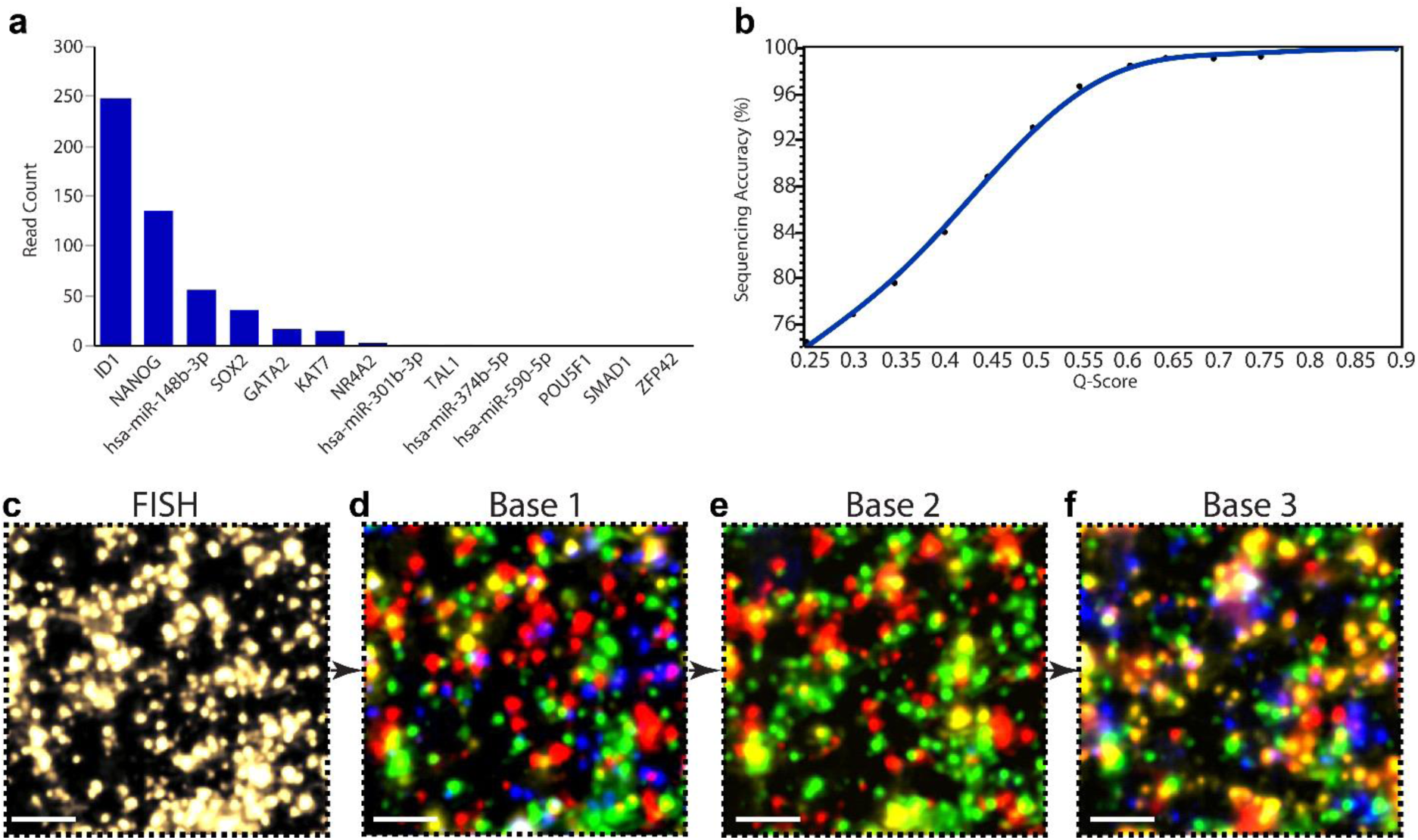
Multiplexed in-situ RNA detection with BOLORAMIS: a) expression profiling of 14 transcripts consisting of miRNA and mRNA in hiPSC. b) in-situ sequencing accuracy as a function of barcode quality scores. c) in-situ sequencing consists of first imaging all BOLORAMIS amplicons with a FISH probe targeting the sequencing adapter on padlock probe. The FISH probe is stripped and hybridized with a sequencing primer and each base is interrogated with sequence-by-ligation. Base-calling was performed by computational image analysis to extract clusters, and fluorescence intensity measurement across cycles. Scale bar: 5 µm

## Discussion

We have shown BOLORAMIS is a RT-free, in-situ single-molecule RNA detection method capable of visualizing single or multiplex coding and small non-coding RNA targets. This is the first demonstration of simultaneous detection of coding and small non-coding RNA with in-situ sequencing. Each BOLORAMIS barcoded amplicon results from a precise, target-RNA dependent single-molecular ligation event, nearing single-base specificity. BOLORAMIS and random-hexamer FISSEQ are complementary methods with unique, and largely non-overlapping applications. Unlike FISSEQ, BOLORAMIS cannot be used for de-novo discovery applications, since the probe sequences must be pre-determined.

Based on published estimates, SplintR mediated direct RNA detection outperforms random-hexamer-RT FISSEQ for targeted transcriptomics by 2-3 orders of magnitude in detection efficiency (∼0.005% vs ∼20%), and circumvents random-primed RT related artifact and over-representation of abundant housekeeping RNA like rRNA^3–5,19,21^. Further, this approach retains the key advantages of the barcoded padlock probe system (multiplexing and targeting sensitivity), without the need for RT or cost-prohibitively expensive LNA modified primers^6,8^.

Due to the short footprint needed for RNA-splinted DNA ligation, BOLORAMIS enables access to transcripts ∼2 orders or shorter than tiled smFISH based methods (∼20 nt vs ∼1500 nt, respectively), opening a vast repertoire of previously inaccessible RNA species for both single or multiplex analysis, including small non-coding RNA, which have traditionally be challenging to detect^29,30^. To the best of our knowledge, this is also the first reported direct comparison of bulk miRNA measurements with in-situ single-molecule miRNA measurements.

### Cost analysis

We estimate BOLORAMIS assay to be over an order or magnitude cheaper than LNA based assays, and within the cost of typical high-content screening (∼1.5$/well) (Supplemental table 8). BOLORAMIS requires far fewer probes than smFISH based approaches. We expect the cost to reduce another 1-3 orders of magnitude with multiplexing. BOLORAMIS library construction workflow is straightforward, robust and can be completed within 1-2 days and amenable to automation.

### Applications towards human cell atlas

Human Cell Atlas (HCA) aims to generate a comprehensive reference maps of all human cells. While single-cell RNAseq (scRNA-seq) methods generate the highest sequencing depth, lose spatial information, and scale more slowly. We believe BOLORAMIS can be a highly complementary approach for spatially mapping transcriptomic signatures obtained from bulk/scRNA-seq methods to identify new cell-types in a high-resolution spatial context. Further, BOLORAMIS may be valuable in constructing large-scale spatial maps of RNA splice variants non-coding RNA.

## Limitations

Success of any targeted RNA detection method, including BOLORAMIS, depends on probe design, and can vary in efficiency based on RNA availability, probe hybridization T^m^ and ligation junction parameters. We anticipate that improvements in probe design algorithms might directly yield improvements in detection efficiency. While PBCV1 based direct RNA detection offers a high detection specificity, the overall fidelity needs to be further improved for applications pertaining to single nucleotide polymorphism detection, without compromising sensitivity^31^. We are actively working on improving In-situ sequencing read depth, read-length and improved base-calling algorithms with error-correction codes. We expect use of partition sequencing in combination with expansion microscopy to vastly overcome a vast majority of the existing current resolution limitations^3,32^.

## Supplemental methods and text

### Cell Culture

Uninduced PGP1 iNGN cells were cultured as described earlier^24^. Briefly, standard tissue culture plates were precoated with 50 µg/ml poly-D-Lysine (PDL) for 1 hour, rinsed in sterile distilled water, and coated with Matrigel hESC-qualified Matrix (354277, BD Biosciences) for one hour at room temperature. The PDL coating was found to help in iPSC attachment. iPSCs were maintained in mTeSR™1 (Stemcell Technologies, Catalog #05850) changed every 24 hours. For passaging, cells were incubated with 1 ml of Gentle Cell Dissociation Reagent (Stemcell Technologies, Catalog#07174) for 8-10 minutes, collected in a 15 ml conical tube with additional mTESR media, and centrifuged at 300g for 5 minutes. Cells were re-plated in fresh mTESR media containing 10µM Y-27632 (ROCK inhibitor) (Stemcell Technologies Catalog# 72302). For high-content RNA profiling, PGP1 hiPSC cells were seeded at 10000 cells/well in 384 well glass bottom plates (Matriplate: MGB101-1-2-LG-L), Media was replaced with fresh mTESR after 24 hours, and allowed to culture for 24 hours prior to fixation. HeLa cells were cultured to confluency, and treated with 0.05% Trypsin-EDTA (Life Technologies), re-suspended in 1X DMEM (Invitrogen) and seeded on six channel flowcells (μ-Slide VI0.4 ibiTreat, Ibidi) and cultured for 24-48 hours prior to fixation.

### Tissue processing

All procedures involving animals are approved by the Harvard University Institutional Animal Care and Use Committee. Mice (adult, female C57BL/6) were and perfused transcardially with 4% paraformaldehyde in 1 x PBS. Brains were dissected out, left in 4% paraformaldehyde at 4C for one day, and then sunk in 30% sucrose for one day. Brains were then embedded in Optimal Cutting Temperature (OCT) matrix and sliced on a cryostat at a thickness of 14 um. Slices were collected on charged glass microscope slides and stored at −20C until use.

Slides were allowed to warm to room temperature and then washed with PBST (PBS + 0.5% Tween-20). A solution of pepsin (2 mg/mL in 100 mM HCl) was applied to slides for 3 minutes. After washing with PBST, slices were then washed with 70%, 85% and 100% ethanol (5 minutes each) before incubating in 100% ethanol for one hour at 4 °C. Slices were then warmed to room temperature and washed with PBST immediately before library preparation.

Fresh-frozen human brain tissue sections were washed with 2x PBST (PBS + 0.5% Tween-20), 5min each and 100% ethanol) at 4 °C. Tissues were split in two groups for detection using BOLORAMIS or RT-padlock method. For RT padlock, tissues were treated 3x with PBST, and incubated overnight with 200 µl of RT reaction mix per slide, followed by wash with PBST. RT reaction mix consisted of 5 µM random decamer RT primers, 20 µl of 200U/ µl RevertAid H Minus M-MuLV Reverse transcriptase (Fermentas), 10 µl of 10mM dNTP (Fermentas), 2 µl BSA (20 µg/µl), 5 µl RiboLock RNAse Inhibitor (40U/ µl, Fermentas), 40 µl RT 5X RT buffer and 120 µl water. Padlocks were hybridized for 4-6 hours, rinsed with 1X PBST, and ligated with 200 µl Padlock ligation mix, which consisted of 20 µl of 10x Ampligase buffer, 0.2 µl of 100uM padlock probes, 1 µl of 100U/ µl Ampligase, 1 µl of 10 mM dNTP, 16 µl of 5U/ µl RNase H (Fermentas), 20 µl of 2U/ µl Phusion DNA polymerase 0.2U/ µl Stoffel fragment (Applied Biosystems), 5µl of 40U/ µl RiboLock RNase Inhibitor (Fermentas), 20 µl of 500mM KCl, 40 µl of 20% formamide, 100ul H2O. The reaction was incubated at 37 °C for 30 min, then 45 °C for 45 mins followed by 2x wash with 1X PBST, followed by overnight RCA.

### Cell fixation, permeabilization and probe hybridization for relative sensitivity and TDR calculation

Cells were fixed with 4% paraformaldehyde (Electron Microscopy Sciences) for 15 minutes, rinsed with PBS (Life Technologies) and permeabilized with Triton X-100 (Thermo Scientific) for 20 minutes. 100 nM of each padlock probe along with the RCA primer in Hybridization Buffer (10% formamide in 6X SSC buffer) were added to the sample and incubated for 1 hour at 37 °C, followed by three 5 minute washes with Hybridization Buffer to remove excess unhybridized primer and probe. Ligase mix containing 210 nM SplintR® Ligase (New England Biolabs) in 1X SplintR Ligase Reaction Buffer was added to the sample and incubated for 1 hour at room temperature followed by three rinses with the Hybridization Buffer to remove the unligated products and enzyme. The samples were then incubated with the Rolling Circle Amplification Mix (1U/µL NxGen Phi29 DNA polymerase (Lucigen), 250 pM dNTPs, 200 µg/ml BSA, 40 μM Amino-Allyl dUTP (Invitrogen)) in 1X Phi29 DNA Polymerase Buffer) for 2 hours at 37 °C followed by three 5-minute washes with 2X SSC buffer. Finally, the samples were stained with 500nM Cy3 labelled fluorescent probes in 6X SSC for 30 minutes, washed with 6X SSC to remove unbound fluorescent probes and imaged with a Leica TIRF fluorescence microscope (Leica, Model DM16000B).

### TDR calculation

A set of 11 probes targeting ACTB mRNA was designed such that the 5’ hybridization arm length was systematically varied at 2 base intervals. The total hybridization region was kept constant at 25 nt, and the same ligation junction (dA/A) was maintained for all sets. As a control, another set of 8 probes targeting ACTB was designed with only one base mismatch at the 3’ ligation junction (A>T). Targeted detection was performed as follows: Confluent cultures of HeLa-Cas9 cells (Genecopoeia, SL503) were treated with 0.05% Trypsin-EDTA (Life Technologies), re-suspended in 1X DMEM (Invitrogen) and seeded on six channel flowcells (μ-Slide VI0.4 ibiTreat, Ibidi). Cells were fixed with 4% paraformaldehyde (Electron Microscopy Sciences) for 15 minutes, rinsed with PBS (Life Technologies) and permeabilized with Triton X-100 (Thermo Scientific) for 20 minutes. 100 nM of each padlock probe along with the RCA primer in Hybridization Buffer (10% formamide in 6X SSC buffer) were added to the sample and incubated for 1 hour at 37 °C, followed by three 5-minute washes with Hybridization Buffer to remove excess un-hybridized oligos. Ligase mix containing 210 nM SplintR® Ligase (New England Biolabs) in 1X SplintR Ligase Reaction Buffer was added to the sample and incubated for 1-2 hours at room temperature followed by three rinses with the Hybridization Buffer to remove the unligated products and enzyme. The samples were then incubated with the Rolling Circle Amplification Mix (1U/µL NxGen Phi29 DNA polymerase (Lucigen), 250 pM dNTPs, 200 µg/ml BSA, 40 μM Amino-Allyl dUTP (Invitrogen)) in 1X Phi29 DNA Polymerase Buffer) for 2 hours at 37 °C followed by three 5-minute washes with 2X SSC buffer. Finally, the samples were stained with 500nM Cy3 labelled fluorescent probes in 6X SSC for 30 minutes, washed with 6X SSC to remove unbound fluorescent probes and imaged with a Leica TIRF fluorescence microscope (Leica, Model DM16000B).

True Discovery Rate (TDR) is defined as the percentage of total signal observed from a perfectly matched probe, *(M/(M+MM)) X 100*. For example, if matched probe results in a mean count of 99 spots/cell, and a mismatched probe results in a mean count of 1 spot/cell, then the TDR is 99%.

**Supplemental Table 1:** BOLORAMIS probe design optimization

| Hybridization Arm length (5') | Hybridization Arm Length (3') | Mean(Mean RCP/Cell Match (TP)) | Sum(Mean RCP/Cell Mismatch (FP)) | Stdev.(Mean RCP/Cell Match (TP)) | Stdev.(Mean RCP/Cell Mismatch (FP)) | Mean TDR (TP/TP+FP) | Stdev. TDR (TP/TP+FP) |
| --- | --- | --- | --- | --- | --- | --- | --- |
| 2 | 23 | 0.214219 | 0.270981 | 0.100637 | 0.069789 | 70.76881 | 13.89577 |
| 4 | 21 | 0.033471 | 0.145825 | 0.003885 | 0.005786 | 40.83457 | 4.611592 |
| 6 | 19 | 2.93611 | 1.418978 | 1.889627 | 0.173507 | 84.02173 | 6.058464 |
| 8 | 17 | 24.85054 | 3.78125 | 2.663579 | 0.201652 | 95.09923 | 1.197424 |
| 10 | 15 | 32.81242 | 4.710869 | 3.88461 | 0.234999 | 95.38814 | 0.904629 |
| 12 | 13 | 60.99675 | 7.24775 | 5.252832 | 0.055003 | 96.17407 | 0.30573 |
| 14 | 11 | 64.6397 | 12.38749 | 3.939773 | 0.3411 | 93.97113 | 0.750441 |
| 16 | 9 | 66.44758 | 22.28406 | 7.863359 | 1.196514 | 89.81306 | 2.398746 |
| 18 | 7 | 75.85146 | 7.864624 | 14.45101 | 0.424285 | 96.64024 | 0.279581 |
| 20 | 5 | 3.230126 | 0.062561 | 1.04143 | 0.018086 | 99.21563 | 0.695311 |
| 22 | 3 | 0.024607 | 0.063314 | 0.00577 | 0.018452 | 63.20762 | 32.61173 |

### BOLORAMIS probe Design

All probes were 60nt or less, with an 18-25 nt RNA targeting region and a 35 nt barcoded linker. The barcoded linker sequence consisted of a universal 23 nt sequencing anchor with built-in secondary structure, flanked by a 6 base HD-4 barcode at either side for multiplexing. The sequencing anchor was used for FISH and in-situ sequencing. Phosphorylated probes were ordered from IDT in 10 nanomoles scale, at a stock concentration of 100 µM in plate format, allowing rapid design-build test cycles.

### Target Selection

77 TFs spanning a broad expression range (log2 FPKM 0-9) were mined from PGP1 iNGN bulk-RNAseq data^24^. A total of 192 probes targeting 77 independent mRNA (TFs) were designed with up to 5 independent probes targeting independent region per RNA. Targeting sequences with high minimal annotated cross-hybridization were mined from Affymetrix Human Primeview arrays (Affymetrix). For all mRNA probes, total targeting region was kept constant at 25 bases, and probes were designed with asymmetric hybridization arm lengths (18/7). Ligation junctions were distributed normally around the expected frequency (mean 12 probes/Ligation Junction +/-Stdev.) with the exception of junction CC (n=4, ∼2%).

All targeting regions were 25 nt or less, and total probe length was 60 nt or less. All probes targeting mRNA, MIP capture sequence was kept constant at 25 nt, with an asymmetric ligation junction positioned 18 nt from the first base of 5’ targeting RNA sequence. For miRNA capture probes, since the total available capture length was limited, ligation junctions were selected between 7 and 15 nt and selected against G in donor/acceptor positions (Since SplintR has been reported to be inhibited by G in either donor or acceptor positions). The mean miRNA probe melting temperature was 64.8 +/-Stdev. of 4.46 (n=77). The mean melting temperature for mRNA targeting probes was 74.35 with a Stdev. of 5.27 °C (n=192) ^33^.

### miRNA probe selection

77 probes targeting 77 mature human miRNA sequences were designed. miRNA targets were selected, guided from previously reported miRNA expression values in human PGP1 iPSC. Mature human miRNA sequences were mined from miRbase. For miR with more than one mature sequences, the mature sequence with higher reported RNA seq values were selected. The mean miRNA length was 22 nt with a Stdev. of +/-0.93. The smallest mature miRNA target sequence was 18 nt (hsa-miR-151b). Probe-ligation junction positions were determined between positions 7-15 nt from 5’, with a mean position of 11 +/-2.6 nt (Stdev.). Positions were selected against G in donor or acceptor positions to increase ligation efficiency (Supplemental Table 2)^19,23^. The mean overall miRNA targeting probe length was 57.2 nt long (+/-.93 Stdev.), with minimum and maximum lengths being 53 and 59 nts respectively.

**Supplemental Table 2:** miRNA Probe ligation junction composition

| **Donor/Acceptor** | **n** |
| --- | --- |
| AA | 5 |
| AT | 6 |
| CT | 6 |
| GC | 1 |
| TA | 41 |
| TC | 4 |
| TT | 14 |
| **Total** | **77** |

### High-content BOLORAMIS assay

Probes were diluted and used between 0.01 µM to 10 µM final concentration in hybridization buffer consisting of 6X SSC with 10% Formamide. Probes were hybridized for upto 16 hours at 37 °C either on a in a warm room, followed by a brief wash in PBS to remove excess probes. SplintR ligation mix was prepared fresh and added slowly on the samples. Ligation reaction was carried out for 2 hours at room temperature or 37 °C with 1X SplintR Ligase mix. SplintR ligase mix consisted of SplintR ligase at 250 nM final concentration in 1X SplintR Ligase reaction buffer. As reported earlier, we also observed a high rate of non-cellular amplicons with SplintR ligase concentrations above 1 µM^22^. RCA mix consisting of 0.6 U/µl of Phi29 Polymerase (Lucigen, 30221-2) in 1X Phi29 polymerase buffer, 0.25 mM dNTPs, in DEPC treated water was added immediately after ligation step. Cells were incubated at 37 °C for 90 minutes, or overnight.

### RNA profiling, high-content imaging and Image Analysis

Fully automated images were acquired from 3 positions chosen randomly from each well on ImageXpress Micro high content screening microscope (MDS Analytical Technologies) using a 40x magnification. Wells with bright fluorescent artifacts were eliminated from analysis. Images were converted from 16bit to 8bit TIFFS, and analyzed using a fully automated pipeline in CellProfiler^34–36^. On average, 1410 cells were imaged per target RNA, 93% of targets imaged consisted of at least 500 cells/ target or more. For each probe, on average 669 cells were imaged, and ∼96% of probes consisted of at least 250 cells or more. For individual probes, mean sample size consisted of 669 cells/ probe with ∼96% of probes containing at least 250 cells. Images were converted to a Maximum Intensity Projection (MIP). Nuclei were defined by automated thresholding and size range 40-200 pixels using shape descriptors. Cell outlines were calculated using CellMask membrane stain (ThermoFisher) and using nucleus position as “seeds” objects. Cytoplasm area was calculated by subtracting the nuclear area from cell area. Puncta images were enhanced using a top-hat filter to enhance the rolling circle amplicons. Watershed segmentation was used to separate amplicons lying in close proximity. All steps were performed using CellProfiler 2.2.0. 3D rendering of images was done using IMARIS (Bitplane). Images were quality checked individually, and wells with any bright fluorescent artifacts were manually eliminated from analysis.

### Sequencing by ligation

Sequencing by ligation was performed with performed with as described by Ke et al. 2013 with a few modifications^6^. 1) Amplicons were stripped of any detection (FISH) probe, followed by 2) incubation with a sequencing anchor, followed by 3) ligation with a pool of degenerate nonomers with one determined position for interrogation. 4) Once ligated, the samples were washed 3x in 6x SSC, and imaged. 5) The signal was “reset” by stripping the sequencing primer and ligated fluorescent product, followed by repeating steps 2-5 repeatedly until sufficient bases have been imaged. Nonamers were synthesized individually with Integrated DNA Technologies (Newark, NJ).

### Sequencing by Ligation

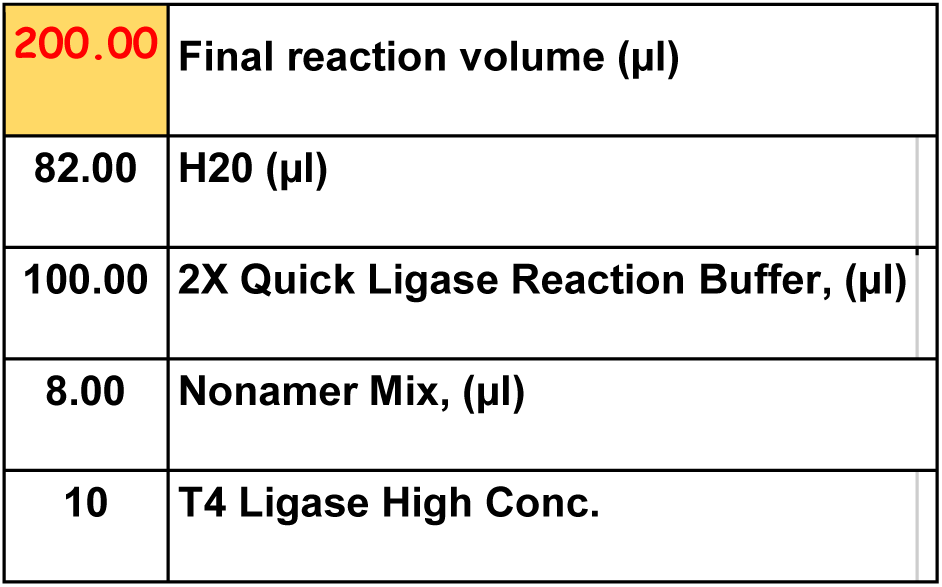

### Image Acquisition, registration and base-calling

Wide-field XYZT image series were obtained on a Zeiss Axio Observer microscope or a Leica inverted DMI6000 TIRF microscope. Focused image-slices were extracted using “Find Focused Slices” plugin (ImageJ) and resulting images were z-projected, labeled and saved as 16bit TIFF images using ImageJ.

Time-series was registered by first performing a Translation, Rotational and Rigid registration in ImageJ using Stackreg plugin^37^. Further elastic-registration was performed to correct for local elastic warping of cells using bUnwarpJ algorithm^38^ (Supplemental Script 1 and 2). Binary-amplicon measurement masks were generated and used to extract amplicon intensities over imaging cycles (Supplemental script 3). A quality score (Q-Score) was calculated for each base as described in Ke. R et al., 2013^6^. In brief, a Q-score was defined as the maximum signal divided by the sum of all channels. For example, if intensities are as follows: A=1000, T=10, C=20, G=30, then Q score for that base = 1000/ (1000+10+20+30) =0.94. The minimum Q-score for all the bases in the barcode were assigned as the overall barcode Q-score. For example, if the barcode is ATC, and the Q scores for each base are A=.9, T=.8, C=.5, then the Q score for the barcode ATC would be 0.5. The quality score can range from 0.25 to 1. The frequency of each barcode sequence was extracted after a Q-score threshold of 0.6.

### Probe Statistics Calculation

Summarized Statistics for each probe was calculated from mean single-cell BOLORAMIS expression values (Puncta Counts/Cell). To estimate dispersion of probability distribution for independent probes targeting the same RNA, COV was calculated for each RNA targets with at least 2 or more probes targeting independent regions.

### Correlation with Bulk Measurements

Mean single-cell BOLORAMIS expression per target were used for correlation analysis. Bulk expression values for PGP1 iNGN cells were mined from Busskamp et al, 2014^24^. To compare BOLORAMIS miRNA measurements, we first identified all miRNAs with published bulk nCounter measurements^24^. Since miRNA precursors produce mature miRNA preferentially from 5’ or 3’ fold-back strand, they are known to accumulate at different rates. As a consequence, a given mature miR ids with a −3p or −5p suffix cannot be compared directly^39^. To be as accurate as possible, we identified all targets from our library with an exactly matched nCounter miRNA measurement form PGP1 iSPC (15/77), which were used for calculating the Pearson’s correlation (below).

**Supplemental Table 3:**
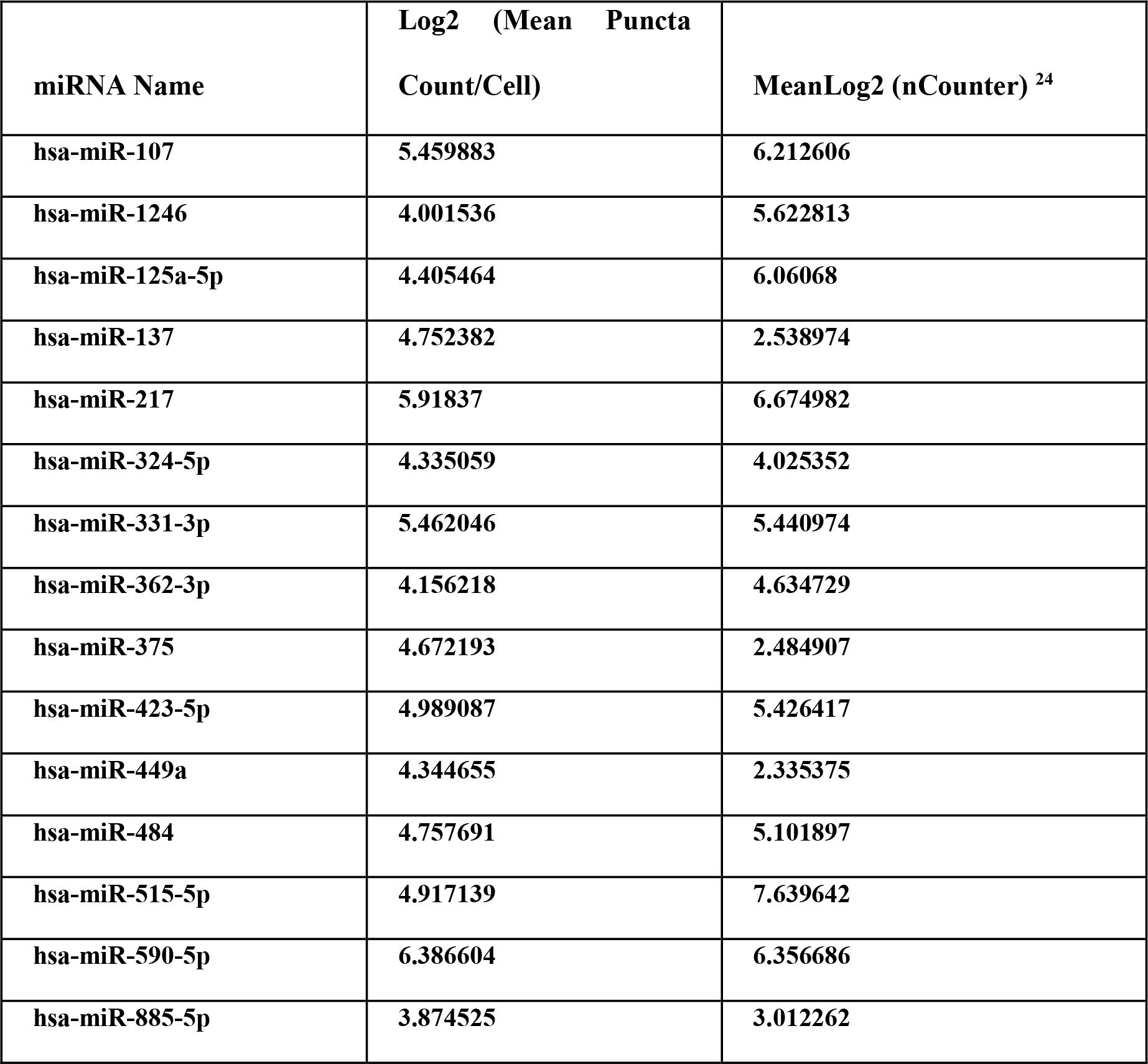
Bulk Mean BOLORAMIS puncta count/cell and nCounter values for miR with available bulk measurements

| **miRNA Name** | **Log2 (Mean Puncta Count/Cell)** | **MeanLog2 (nCounter) <sup>24</sup>** |
| --- | --- | --- |
| hsa-miR-107 | 5.459883 | 6.212606 |
| hsa-miR-1246 | 4.001536 | 5.622813 |
| hsa-miR-125a-5p | 4.405464 | 6.06068 |
| hsa-miR-137 | 4.752382 | 2.538974 |
| hsa-miR-217 | 5.91837 | 6.674982 |
| hsa-miR-324-5p | 4.335059 | 4.025352 |
| hsa-miR-331-3p | 5.462046 | 5.440974 |
| hsa-miR-362-3p | 4.156218 | 4.634729 |
| hsa-miR-375 | 4.672193 | 2.484907 |
| hsa-miR-423-5p | 4.989087 | 5.426417 |
| hsa-miR-449a | 4.344655 | 2.335375 |
| hsa-miR-484 | 4.757691 | 5.101897 |
| hsa-miR-515-5p | 4.917139 | 7.639642 |
| hsa-miR-590-5p | 6.386604 | 6.356686 |
| hsa-miR-885-5p | 3.874525 | 3.012262 |

**Supplemental Table 4:** Effect of ligation junction on BOLORAMIS correlation with RNAseq

| **18/7 Junction** | **r (Pearson)** | **R<sup>2</sup>** | **N (number of probes per category)** |
| --- | --- | --- | --- |
| **TC** | **0.928** | **0.861** | **10** |
| **CC** | **0.856** | **0.732** | **4** |
| **AC** | **0.573** | **0.329** | **13** |
| **CA** | **0.57** | **0.325** | **14** |
| **GG** | **0.395** | **0.156** | **15** |
| **AT** | **0.37** | **0.137** | **12** |
| **AG** | **0.166** | **0.027** | **10** |
| **TA** | **0.09** | **0.008** | **10** |
| **GC** | **-0.032** | **0.001** | **8** |
| **TT** | **-0.045** | **0.002** | **11** |
| **CT** | **-0.108** | **0.012** | **13** |
| **GT** | **-0.131** | **0.017** | **19** |
| **AA** | **-0.203** | **0.041** | **9** |
| **GA** | **-0.225** | **0.05** | **19** |
| **CG** | **-0.275** | **0.076** | **9** |
| **TG** | **-0.351** | **0.124** | **16** |

192 probes targeting mRNA were separated by ligation junction composition, and Pearson correlation coefficients with bulk RNAseq FPKM values were calculated independently for each group^24^.

**Supplemental table 5:** Effect of probe hybridization temperature on BOLORAMIS correlation with published RNAseq in hiPSC

| **T<sub>m</sub>** | **Pearson Correlation (r) Mean Puncta**<br>**Count/Cell vs Bulk RNA-seq FPKM** |
| --- | --- |
| **62-70** | **-0.202** |
| **70-74** | **0.202** |
| **74-76** | **-0.029** |
| **76-80** | **0.191** |
| **80-82** | **0.227** |
| **82-86** | **0.553** |

BOLORAMIS Count/cell were correlated with published bulk RNA-seq FPKM values^24^. Probes with higher hybridization temperatures correlated better with bulk FPKM values than lower ones on average. Probes with highest T^m^. Fell in the range of 82-86 displayed a higher Pearson’s correlation (r=0.533) than comparison to probes with Tm in the range of 62-70, 70-74, 74-76, 76-80, 80-82, 82-86.

Barcoded BOLORAMIS padlock probes can be ordered ready-to-use from commercial vendor (IDT), and offers a significant advantage for rapid testing of hypothesis by saving oligo chip-synthesis, oligo amplification, purification and phosphorylation costs. BOLORAMIS RNA targeting region is shorter than most reported MIPs for in-situ RNA capture^40^. Shot oligo design allows cheap assays with and quick turnaround time and minimizes synthesis errors. We routinely order ready to use phosphorylated probes in 96/384 well formats at 100 µM concentration (IDT, ∼$5.2 per probe at 7c/base + $1 phosphorylation cost, 2-4 days turnaround time for ready to use BOLORAMIS probes. Total assay-time 1-2 days depending on the incubation steps.

**Supplemental table 6:** Effects of elastic registration on in-situ sequencing

|  | **Without**<br>**elastic**<br>**registration** | **With**<br>**Elastic**<br>**registration** |
| --- | --- | --- |
| **Total puncta** | **1285** | **520** |
| **Exact match** | **1134** | **511** |
| **% match** | **88.24903** | **98.26923** |
| **% of input Barcodes identified** | 85.71429 | 64.28571 |
| **Number of barcode library match**<br>**(library=14)** | 12 | 9 |
| **Total Unique Barcodes Identified** | **37** | **14** |
| **Quality Filter Cutoff** | **>=0.6** | **>=0.6** |

**Supplemental table 7:** Multiplexed in-situ sequencing of coding and small non-coding RNA

| **Gene Name** | **Expected**<br>**Barcode** | **Single-plex**<br>**puncta/cell** | **V5-**<br>**Multiplex** | **V6-**<br>**Multiplex** |
| --- | --- | --- | --- | --- |
| ZFP42 | ACG | 236.9585 | 5 | 0 |
| GATA2 | CAT | 231.4655 | 309 | 17 |
| MIMAT0000759 | ATC | 217.824 | 344 | 56 |
| SMAD1 | GTG | 209.1381 | 11 | 0 |
| ID1 | AAA | 205.3245 | 929 | 247 |
| KAT7 | TCA | 181.3808 | 76 | 15 |
| POU5F1 | CGC | 113.5869 | 1 | 0 |
| MIMAT0004958 | TAC | 92.18129 | 9 | 1 |
| MIMAT0003258 | AGG | 83.66802 | 4 | 0 |
| MIMAT0004955 | CTA | 78.5593 | 11 | 0 |
| SOX2 | CAG | 57.68784 | 601 | 36 |
| NANOG | GTT | 56.3425 | 586 | 135 |
| NR4A2 | GGA | 32.57613 | 21 | 3 |
| TAL1 | TCC | 27.80296 | 0 | 1 |

**Supplemental table 8:** BOLORAMIS Cost/ reaction

|  | Cost Per unit | Supplied as | Unit Definition | Amount used per 25µl reaction | Cost Per Reaction | Notes |
| --- | --- | --- | --- | --- | --- | --- |
| Phosphorylated BOLORAMIS Padlock probes | 5.2 | 10 | nm, supplied at 100 µM concentration | 0.125 | 0.065 | Ordered directly from IDT |
| SplintR Ligase | 327 | 6250 | Units, supplied at 25U/ul | 15 | 0.7848 |  |
| Phi29 Polymerase | 419 | 10000 | Units, supplied at 10U/ul | 1.5 | 0.06285 |  |
| Amino-Allyl dUTP |  |  |  |  |  |  |
| Detection Probe | 200 | 25 | nM | 0.1 | 0.2 | Cost calculated for a 23nt CY3 labeled FISH probe, ordered at 250 nmole scale with HPLC purification from IDT. Typically we receive 1/10th of the ordered scale (25nM). 25nM = 250 µl of 100X stock probes, which is sufficient for 1000, 25 µl reactions. Total cost per reaction would be 200\$/1000 reactions=0.2\$/reaction |
| Miscellaneous (PBS, water etc) |  |  |  |  | 0.25 |  |
| dNTP cost not included |  |  |  | Total cost/ 25 µl assay | 1.36265 |  |

BOLORAMIS assays were calculated for a 25 μl assay volume and includes the cost of oligonucleotide probe synthesis and enzymes. LNA probe calculation was performed from available costs from a commercial vendor (Exiqon), calculated at $54.96 per assay of equivalent volume, at the suggested final concentration of 50nM^29^. Overall, BOLORAMIS allows one to screen and visualize large number of RNA with minimal starting costs. For example, the initial investment to large scale miRNA detection is ∼33 fold cheaper in comparison to LNA probes ($33,858.44 initial investment to obtain 77 LNA-FAM labeled probes at $439.72 per probe versus. In contrast, all BOLORAMIS reagents (probes, enzymes and FISH probes) for detecting an equivalent number of miRNA can be purchased for ∼$1000.

## Acknowledgements

We thank Thouis Ray Jones for helping with software development for in-situ sequencing. We also gratefully acknowledge Jay H. Lee, Evan Daugharty, Brian Turczyk, Rich Terry, Samuel Inverso for Rigel Chan, Elaine Lim, Dima Ter-Ovanesyan, Alexander Shineman Garruss and Alejandro Chavez for engaging discussion, advice and insights on various steps. We also thank Daniel Collett (IDT) for support on oligo probe synthesis. We gratefully acknowledge funding and support from The Center for Genomically Engineered Organs (CGEO, RM1HG008525, and NHGRI), Department of Energy (DOE, DE-FG02-02ER63445), NIH 1R01MH113279-01 (NIH, G.C. and B.A.Y.) and The Wyss Institute for Biologically Inspired Engineering at Harvard University.

## Author Contributions

EPRI initiated the project and made high-level designs and plans with GMC and JA. EPRI, SP, SL and KJ planned and executed experiments, EPRI and MF developed image-processing and data analysis pipelines. EPRI, SP, SL collected, analyzed and interpreted data. SP and KJ performed probe optimization experiments. SL performed high-resolution confocal imaging. JM mined and helped design splice-junction experiments. TF helped with custom, sophisticated microscopy setup (for in-situ sequencing and high-resolution imaging). SR helped obtain high-content imaging technical setup. EPRI performed high-content imaging, automated quantification and statistical analysis. REK, SP, EPRI designed and executed mouse brain tissue experiments. ATW, DG, FC, SA and AS and ESB shared unpublished data and insights, and helped plan experiments. GMC, JA, DG, SA helped troubleshoot detection, in-situ sequencing data analysis. BY, LA provided human brain tissue samples, advice and insights. DM helped with RT-padlock experiments. CC explored early strategies for in-situ RNA detection using in-situ PCR. PR explored early padlock detection methods. AS, TF, KJ, MF and EPRI planned and developed in-situ sequencing automation strategies. GMC supervised the project. EPRI wrote the manuscript with GMC and JA, with contributions, edits and revisions from all authors.

## Supplemental Figures

**Figure S1:**
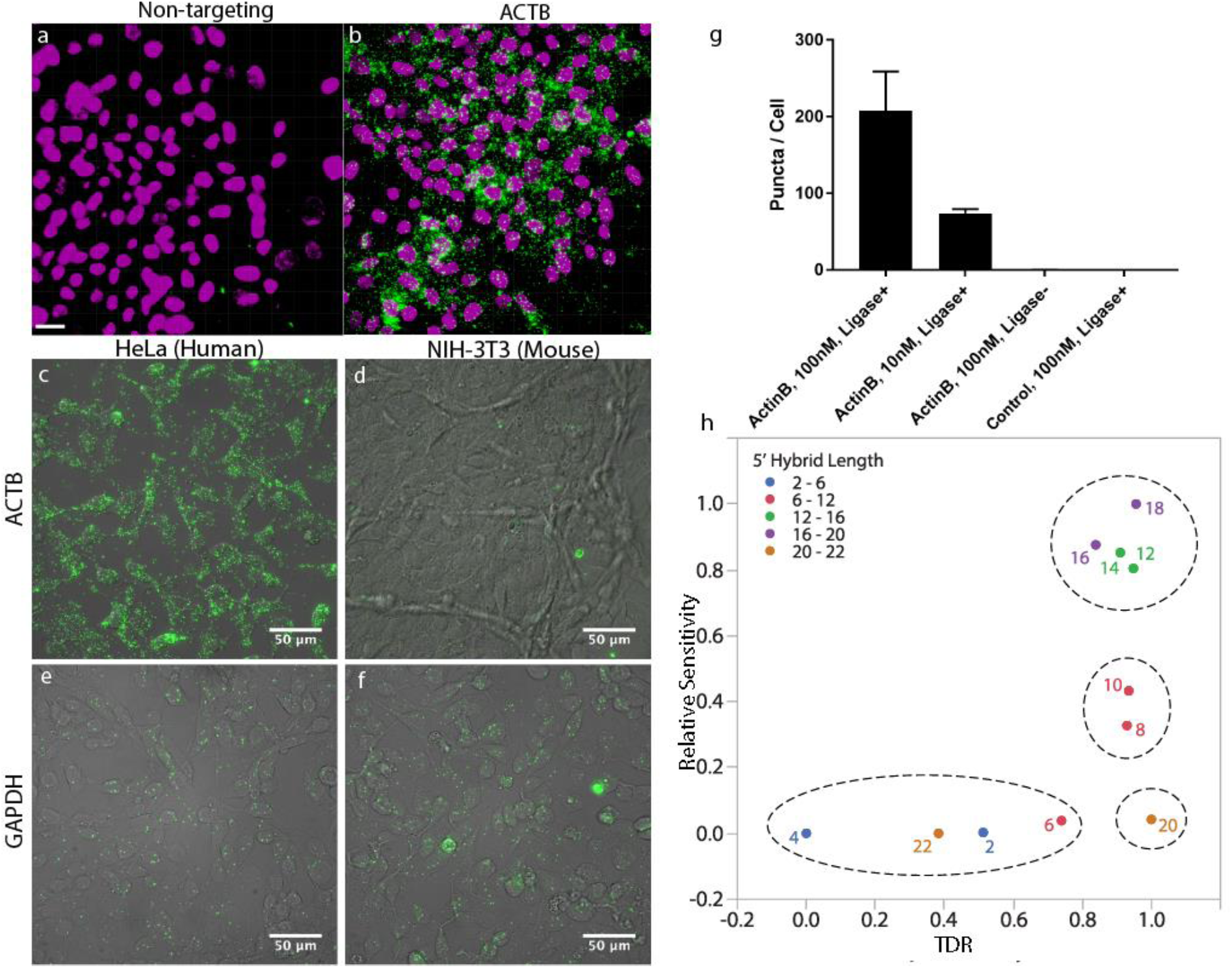
BOLORAMIS detects RNA with high specificity: Non-targeting (negative control) probes result in almost no signal (a), in comparison to probes targeting ACTB (b). Using probes designed against human-specific 3’ UTR resulted in overwhelming signal from HeLa cells (c), but not Mouse NIH 3T3 cells (d). In contrast, probe designed against a conserved 3’UTR resulted in dense punctate signal from both HeLa (e) and NIH 3T3 cells (f). (g) Results of image quantification indicates highly specific detection of targets. Non-targeting probes (negative controls), or absence of ligase resulted in almost no signal from images. h) Relative sensitivity vs TDR plot for BOLORAMIS probes with varying 5’ DNA: RNA hybrid length results in four unique clusters. Probes with 2, 4, 6 and 22 nt long 5’ ligation junctions had the least TDR and relative sensitivity, likely due to insufficient hybridization length at either 3’ or 5’ termini. Probes with 12, 14, 16 and 18 nt long 5’ termini displayed the highest relative sensitivity and TDR.

**Figure S2:**
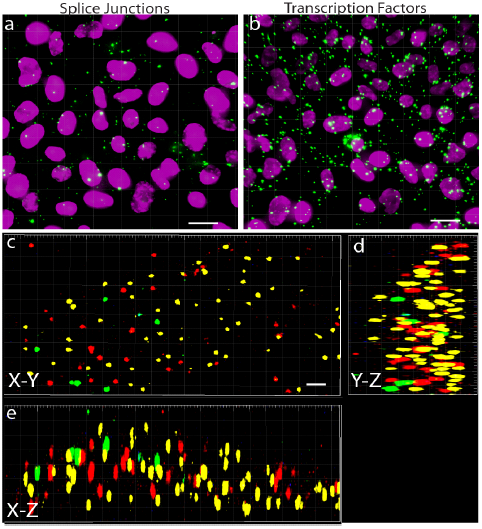
multiplexed in-situ sequencing libraries generated in human cells targeting TFs and Splice Junctions: Pooled BOLORAMIS probes targeting splice-junctions (18-plex, a), and transcription factors (77-plex, b) were used to generate targeted in-situ libraries from HeLa cells. DAPI staining reveals the nucleus, pseudocolored to magenta for clarity, and green dots are BOLORAMIS puncta. (c,d) 3-d rendering of a single HeLa cell showing spatially defined clonal BOLORAMIS amplicons. BOLORAMIS amplicons appear elongated by ∼2 fold in Z-axis due to over-sampling of Z-slices during imaging. Scale-bars 20 μm (a,b), 3 μm (c-e).

**Figure S3:**
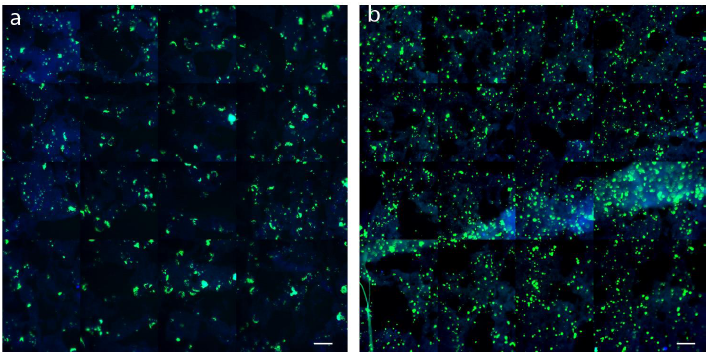
BOLORAMIS and RT-padlock probe on frozen human brain sections: Fresh frozen human brain tissue from a young individual was fixed, permeabilized. The tissues were either processed either by (a) first reverse transcribing the RNA using random decamer followed by padlock probe hybridization, or (b) direct padlock probe hybridization to RNA targeting the identical sequence followed by ligation and RCA. Rolling circle amplicons were visualized using a FISH probe targeting the universal anchor (green puncta). A 1.1 mm2 area of tissue was imaged for each condition. The punctate spots were counted by global thresholding and using analyze-particles feature in ImageJ using the same settings. The punctate spots were normalized over the total area covered by DAPI signal (blue). Overall, BOLORAMIS signal was comparable with RT-padlock probe method (in this particular example, BOLORAMIS generated 2.325 fold higher spots than padlock-probe detection of reverse-transcribed cDNA). Scale bar 50 μm.

**Figure S4:**
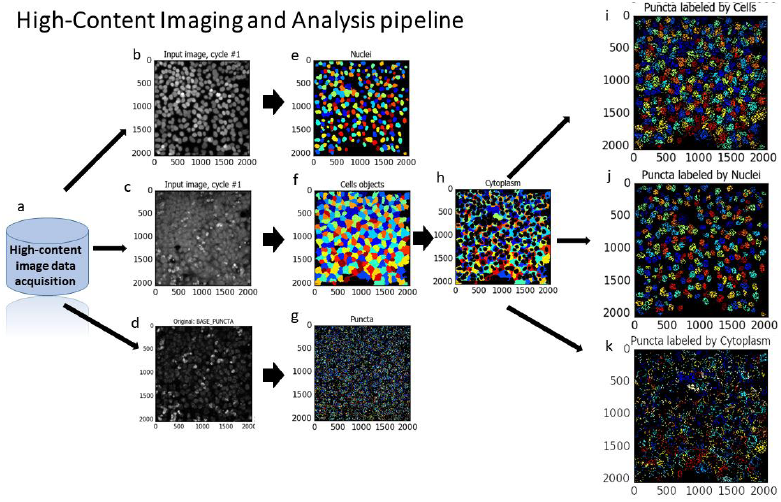
Automated image analysis pipeline: Cells were cultured and processed in parallel in a multi-well plate and stained. a) Thousands of images representing tens of thousands of cells were collected using multiple wavelengths using a fully automated high-content screening microscope and stored in a central database. Z-stacks were projected using Maximum-intensity projection, and Images are down-sampled from 16 bit to 8-bit tiff format. (b-k) A custom CellProfiler pipeline was developed for segmenting and analyzing the images. Briefly, the nuclei are defined by automated thresholding and using shape descriptors (b, e). The cell-outlines are calculated using cell-mask membrane stain and using nucleus as a “seed” objects for each cell (c, f). Individual spots, representing an RNA molecule were detected using a top-hat filter (d, g). Cytoplasmic boundaries are calculated by subtracting nuclear area from total cell-boundaries (h). Finally, the identified puncta and other objects are computationally associated with each cell such that the spatial identity is preserved (i, j, k).

**Figure S5:**
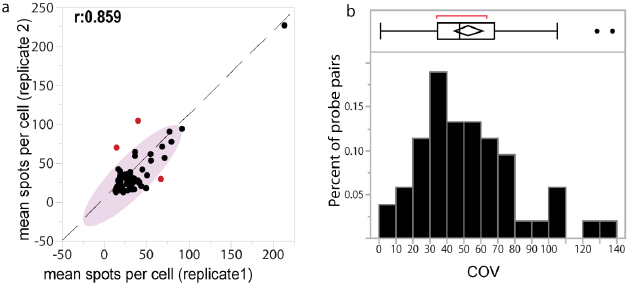
BOLORAMIS Assay reproducibility: Mean single-cell BOLORAMIS values for 77 miRNA’s assayed in replicates are plotted. (a) The Pearson’s correlation coefficient was 0.859 without outlier rejection. (b) Distribution of COV for independent probes targeting the same RNA transcripts is shown.

**Figure S6:**
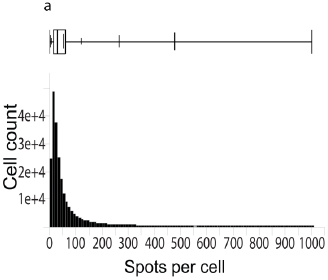
Single Cell distribution of BOLORAMIS spots in human iPSC: Distribution of BOLORAMIS single-cell expression from 217,206 single human iPSC cells. Boxplot on top displays the 25th, 50th, 75th, 95th and 99th quantiles.

**Figure S7:**
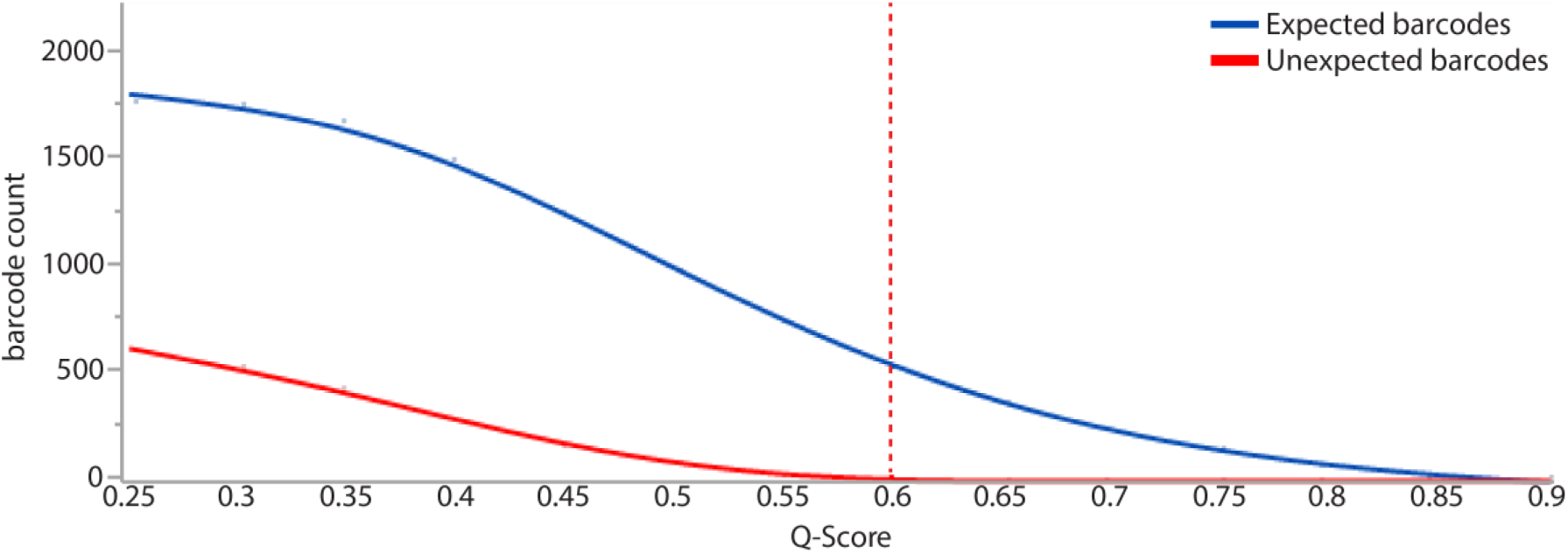
In-situ sequencing barcode frequency vs quality: Barcode deconvolution results in some sequences that do not map to the input barcode library. However calculating a quality filter, and thresholding it results in barcodes with <2% unexpected sequences.

**Figure S8:**
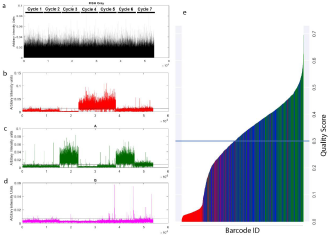
In-situ sequencing of BOLORAMIS: BOLORAMIS generates bright sequenced amplicons with a high signal/noise. Six bases in the minus direction and one base in the plus direction (total of 7 bases) were sequenced with sequence by ligation as reported earlier6. To minimize signal crosstalk between Cy3 and Texas-Red, we eliminated Texas-Red and instead used a 3 color + “blank color” scheme as follows: T: Cy5 (Red plot), A: Cy3 (Green Plot), G: FAM (Magenta plot) and C is blank. In the above example, probes targeting ACTB with the barcode ‘CCATTAC’ in HeLa cells were sequenced. As expected we observed no peaks in the first and second cycles, which correspond to C (blank-color). At cycle 3 and 6 we detected strong signals from Cy3 channel representing the base A (plot b). At cycle 4 and 5, signal was observed only in Cy5 (plot b). As negative control, the barcode was designed without a ‘G’, and as expected the FAM channel (plot d) displayed signal at baseline levels. Quality scores were assigned to each base based on its overall intensity value. Sequenced barcodes with perfect match to expected barcodes are shown in green. Deviation from expected probes by 1 or 2 distances is depicted by blue and red colors. Overall, 95.97% of identified barcodes mapped to the expected barcode within 1 hamming distance (N=8464 amplicons, N post quality filter = 4792, perfect match=38.19%, <=1 Hamming distance=74.65%).

